# A Machine Learning Approach for Long-Term Prognosis of Bladder Cancer based on Clinical and Molecular Features

**DOI:** 10.1101/557470

**Authors:** Qingyuan Song, John D. Seigne, Alan R. Schned, Karl T. Kelsey, Margaret R. Karagas, Saeed Hassanpour

## Abstract

Improving the consistency and reproducibility of bladder cancer prognoses necessitates the development of accurate, predictive prognostic models. Current methods of determining the prognosis of bladder cancer patients rely on manual decision-making, including factors with high intra- and inter-observer variability, such as tumor grade. To advance the long-term prediction of bladder cancer prognoses, we developed and tested a computational model to predict the 10-year overall survival outcome using population-based bladder cancer data, without considering tumor grade classification. The resulted predictive model demonstrated promising performance using a combination of clinical and molecular features, and was also strongly related to patient overall survival in Cox models. Our study suggests that machine learning methods can provide reliable long-term prognoses for bladder cancer patients, without relying on the less consistent tumor grade. If validated in clinical trials, this automated approach could guide and improve personalized management and treatment for bladder cancer patients.

## Introduction

Urothelial bladder cancer is one of the most common malignancies worldwide, with the estimates of 80,470 new cases and 17,670 deaths in the United States in 2019^1^. Although the incidence rate of bladder cancer has been decreasing in recent years, the disease frequently recurs and its mortality rate has remained unimproved^1^. Thus, the development of accurate, predictive prognostic markers is needed to better personalize healthcare management of this disease and improve patient survival, as patients with poorer prognoses may benefit from more intense follow-up, treatment, and healthcare planning.

Patient prognosis often involves some degree of uncertainty due to the nature of predicting future events. In clinical practice, clinicians make prognostic judgments by juxtaposing patients’ clinical information with their own personal experience and prior knowledge for treating patients. Therefore, manual prognostic judgment can vary greatly among clinicians and institutions^2^. In addition, some prognostic factors can depend upon subjective assessments that lack consistency and reproducibility. For example, the WHO 1973 and WHO/ISUP (World Health Organization/International Society of Urological Pathology) classification are the two most widely used methods for establishing tumor grade, but these methods have relatively high intra- and inter-observer reliability^3, 4^. Robertson et al. reported that the inter-observer agreement among 11 pathologists using the WHO 1973 classification was slight to moderate (κ = 0.19 − 0.44)^5^. A study by Yorukoglu et al. compared the inter-observer agreement among 6 pathologists using both classifications and found only a slightly better but still moderate inter-observer agreement with the WHO/ISUP classification (κ = 0.42 − 0.65) than the WHO 1973 classification (κ = 0.19 − 0.65)^6^. Ooms et al. found a limited intra-observer variability (Spearman rank-order correlations coefficients of 0.50–0.67) using the WHO 1973 classification^7^.

Automated computational methods have demonstrated utility in providing unbiased and reliable guidance in clinical decision-making for bladder cancer patients by exploiting large datasets^7^. Furthermore, machine learning approaches require minimal time and resources. To demonstrate the potential utility of computational models, we designed and tested a fully automated machine learning pipeline to predict 10-year bladder cancer survival based on factors from a population-based dataset. Such models are of potential clinical utility, particularly, because there is currently lack of predictors of long-term survival for the general population of patients with bladder cancer.

## Methods

The overview of the pipeline for data collection, model training, and evaluation is shown in Figure 1.

**Figure 1.**
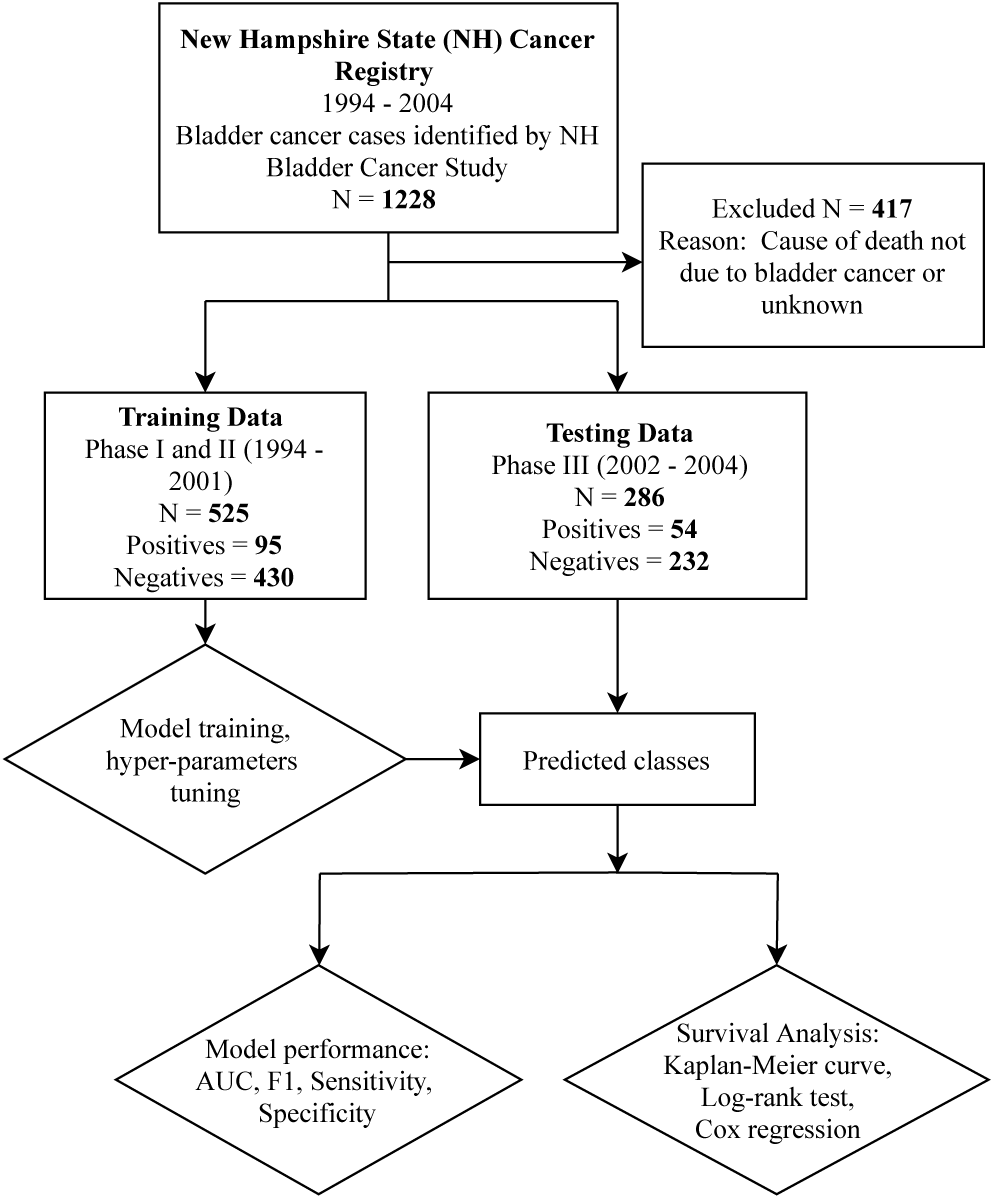
An overview of an automated machine learning pipeline for bladder cancer 10-year survival prediction. A total of 811 bladder cancer patients were split into training and testing datasets. Positive samples included patients with disease-specific deaths within 10 years from the initial diagnosis, and negative samples included patients with a survival of more than 10 years from the initial diagnosis. The model was trained on data from Phases I and II combined and evaluated on data from Phase III. The model’s predictions were compared to histological classification through survival analysis.

### Dataset

Our study utilized 1,228 previously collected histologically confirmed bladder cancer cases from the New Hampshire Bladder Cancer Study that were identified from the New Hampshire State Cancer Registry^9^. The Dartmouth Institutional Review Board approved this study and the use of human patient data in this project with a waiver of informed consent. The dataset consists of cases from three study phases: Phase I included 448 patients diagnosed from July 1, 1994 to June 30, 1998^10^; Phase II included 385 patients diagnosed from July 1, 1998 to December 31, 2001^10^; Phase III included 396 patients diagnosed from July 1, 2002 to December 31, 2004^11^. Cases were restricted to patients aged 25 to 74 years at initial diagnosis in the first two study phases and 31 to 79 years at initial diagnosis in the third study phase. Demographic and risk factor information in this registry was obtained through personal interviews. The high-risk occupations of patients for this dataset were previously described in Colt et al.^12^. Histological slides were requested from the pathology laboratory of initial diagnosis and were reviewed by a single pathologist masked from the submitted diagnosis. Tumor grade according to both WHO 1973^13^ and 1998 WHO/ISUP^14^ criteria were assessed by a standardized re-review of the original histopathology specimens. Tumor stage was reported using the 2002 revised TNM criteria of the American Joint Commission on Cancer^15^. Immunohistochemical (IHC) staining and mutation analysis of molecular biomarkers were previously described in Kelsey et al.^10^. The distribution of our data is similar to that of a larger population-based dataset from the Surveillance, Epidemiology, and End Results (SEER) database^16^.

### Outcome Status

All patients who survived from Phases I and II had follow-ups for at least 120 months. The survival status of patients was determined by examining the Social Security Administration Death Master File^17^. Survival time was calculated from the date of diagnosis to the date of death for patients who did not survive, or to the date when the Death Master File was queried for patients who survived. Patients were labeled as positives and negatives based on whether or not they survived at least 120 months after the initial diagnosis. Patients in Phase III of the study were classified according to their death status at the last follow-up, which was approximately 10 years (129.0±8.90 months) for patients who survived. To avoid misclassification, we removed patients whose causes of death were unknown or not contributed to by bladder cancer. The causes of patients’ death were reported by the National Death Index (NDI). The primary and secondary causes of death for each patient were coded using the International Classification of Diseases (ICD) system. Bladder cancer was decided as a contributor to a patient’s death if it was among one of the primary and secondary causes of deaths according to the ICD codes. As a result, of the 1,228 bladder cancer patients, 417 were removed due to non-bladder-cancer-related deaths or unknown causes of deaths. The remaining patients in this study included 149 persons with disease-specific deaths within 10 years after the initial diagnosis, labeled as positives, and 662 persons who survived more than 10 years, labeled as negatives. To add to the generalizability of our model and follow the chronological order of the collected dataset, we combined data from earlier phases (Phases I and II) into training data and used Phase III as a testing dataset. The ratios between the number of positives and negatives are consistent in both training and testing datasets. The baseline characteristics of patients included in this study are shown in Table 1.

**Table 1.** Characteristics of 811 bladder cancer patients used in this study.

| Variable | Type | Overall | Training | Testing |
| --- | --- | --- | --- | --- |
| n |  | 811 | 525 | 286 |
| Age (SD) |  | 61.17 (10.33) | 59.82 (10.04) | 63.67 (10.41) |
| Sex (%) | Men | 586 (72.3) | 385 (73.3) | 201 (70.3) |
|  | Women | 225 (27.7) | 140 (26.7) | 85 (29.2) |
| Body mass index (SD) |  | 27.80 (5.03) | 28.17 (4.71) | 27.69 (5.12) |
| Family history of bladder cancer (%) | Yes | 40 (4.9) | 25 (4.8) | 15 (5.2) |
|  | No | 724 (89.3) | 453 (86.3) | 270 (94.8) |
|  | Unknown | 47 (5.8) | 47 (9.0) | 0 (0.0) |
| High risk occupation (%) | Yes | 284 (35.0) | 128 (24.4) | 127 (44.6) |
|  | No | 258 (31.8) | 130 (24.8) | 156 (54.7) |
|  | Not available | 269 (33.2) | 267 (50.9) | 2 (0.7) |
| Smoke status (%) | Current | 246 (30.3) | 158 (30.1) | 88 (30.8) |
|  | Former | 397 (49.0) | 252 (48.0) | 145 (50.7) |
|  | Never | 160 (19.7) | 110 (21.0) | 50 (17.5) |
|  | Unknown | 8 (1.0) | 5 (1.0) | 3 (1.0) |
| Smoke pack-years (SD) |  | 40.01 (30.19) | 41.24 (31.06) | 37.88 (28.58) |
| UCC status (%)* | Confirmed UCC | 729 (89.9) | 456 (86.9) | 273 (95.5) |
|  | Not UCC | 17 (2.1) | 12 (2.3) | 5 (1.7) |
|  | Not available | 65 (8.0) | 57 (10.9) | 8 (2.8) |
| Muscle invasiveness (%) | Yes | 100 (12.3) | 58 (11.0) | 42 (14.7) |
|  | No | 705 (86.9) | 466 (88.8) | 239 (83.6) |
|  | Not available | 6 (0.7) | 1 (0.2) | 5 (1.7) |
| WHO 1973 classification (%) | Grade 1 | 336 (41.4) | 193 (36.8) | 143 (50.0) |
|  | Grade 2 | 166 (20.5) | 110 (21.0) | 56 (19.6) |
|  | Grade 3 | 161 (19.9) | 85 (16.2) | 76 (26.6) |
|  | Grade 4 | 65 (8.0) | 61 (11.6) | 4 (1.4) |
|  | Pathology material not reviewed | 17 (2.1) | 17 (3.2) | 0 (0.0) |
|  | Not available | 66 (8.1) | 59 (11.2) | 7 (2.4) |
| WHO/ISUP classification (%)* | CIS | 28 (3.5) | 15 (2.9) | 13 (4.5) |
|  | Papilloma | 2 (0.2) | 2 (0.4) | 0 (0.0) |
|  | PUNLMP | 201 (24.8) | 122 (23.2) | 79 (27.6) |
|  | LG-PUC | 225 (27.7) | 148 (28.2) | 77 (26.9) |
|  | HG-PUC | 196 (24.2) | 109 (20.8) | 87 (30.4) |
|  | Non-PUCHG | 53 (6.5) | 36 (6.9) | 17 (5.9) |
|  | Other | 42 (5.2) | 35 (6.7) | 7 (2.4) |
|  | Not available | 64 (7.9) | 58 (11.1) | 6 (2.1) |
| TNM Stage (%) | CIS | 41 (5.1) | 29 (5.5) | 12 (4.2) |
|  | 0 | 541 (66.7) | 347 (66.1) | 194 (67.8) |
|  | 1 | 123 (15.2) | 90 (17.1) | 33 (11.5) |
|  | 2 | 38 (4.7) | 23 (4.4) | 15 (5.2) |
|  | 3 | 22 (2.7) | 14 (2.7) | 8 (2.8) |
|  | 4 | 40 (4.9) | 21 (4.0) | 19 (6.6) |
|  | Not available | 6 (0.7) | 1 (0.2) | 5 (1.7) |
| P53 mutation (%) | Yes | 14 (1.7) | 14 (2.7) | 0 (0.0) |
|  | No | 186 (22.9) | 186 (35.4) | 0 (0.0) |
|  | Not available | 611 (75.3) | 325 (61.9) | 285 (100.0) |
| P53 intensity (%) | 3+ | 490 (60.4) | 298 (56.8) | 192 (67.1) |
|  | < 3 | 182 (22.4) | 120 (22.9) | 62 (21.7) |
|  | Not available | 139 (17.1) | 107 (20.4) | 32 (11.2) |
| P53 positivity (%) | 50% + | 323 (39.8) | 232 (44.2) | 91 (31.8) |
|  | < 50% | 322 (39.7) | 159 (30.3) | 163 (57.0) |
|  | Not available | 166 (20.5) | 134 (25.5) | 32 (11.2) |
| PTCH LOH positivity (%) | Yes | 119 (14.7) | 119 (22.7) | 0 (0.0) |
|  | No | 45 (5.5) | 45 (8.6) | 0 (0.0) |
|  | Not available | 647 (79.8) | 361 (68.8) | 285 (100.0) |
| First course of therapy (%) | No treatment | 39 (4.8) | 36 (6.9) | 3 (1.0) |
|  | TUR only | 577 (71.1) | 368 (70.1) | 209 (73.1) |
|  | Intravesical BCG* | 96 (11.8) | 63 (12.0) | 33 (11.5) |
|  | Intravesical Chemotherapy | 8 (1.0) | 5 (1.0) | 3 (1.0) |
|  | Radiation + Chemotherapy | 13 (1.6) | 6 (1.1) | 7 (2.4) |
|  | Cystectomy | 78 (9.6) | 47 (9.0) | 31 (10.9) |
| Mean survival time in months (SD) |  | 143.94 (55.94) | 159.89 (58.68) | 114.65 (35.11) |
| Death status at 10-year mark(%) | Dead | 149 (18.4) | 95 (22.1) | 54 (18.9) |
|  | Alive | 662 (81.6) | 430 (77.9) | 232 (81.4) |

\*UCC – urothelial carcinoma, HG-PUC –high grade papillary urothelial carcinoma, LG-PUC – low grade papillary urothelial carcinoma, PUNLMP – papillary urothelial neoplasm of low malignant potential, CIS – carcinoma *in situ*, BCG – Bacillus Calmette-Guerin

### Missing data imputation

We implemented a data imputation strategy for missing covariate data. Training data and testing data were imputed separately. Binary variables with missing values (i.e., *p53* mutation, *p53* intensity, PTCH LOH positivity, high-risk occupation, family history of cancer) were imputed with 0.5 (midpoint value). Body mass index (BMI) was mean imputed. Missing values for smoke pack-years were imputed with ‘0’ for never smokers, and with the mean pack-years for others.

### Predictive Model

We implemented a logistic regression model with demographic characteristics (age and sex), risk factors (history of cigarette smoking, high risk occupation, BMI, and family history of bladder cancer), clinical information (presence of muscle invasiveness and tumor histology), and molecular features (*p53* mutation, *p53* IHC positivity and *p53* IHC staining intensity; PTCH LOH). The features included were putative or potential predictors of bladder cancer survival and are known to be reliable or objective measures. Model implementation was done using Python 3.6 (Python Software Foundation, Beaverton, OR) and Scikit Learn version 0.19.1^18^. Because the distribution of positive and negative classes in the dataset was highly imbalanced, we weighted each class reciprocal to its prevalence in the training dataset and maximized the weighted log-likelihood loss function to estimate the parameters of the model. The class weights were calculated using equation (1) as suggested by King^19^ so that the minority class was more emphasized in the model:

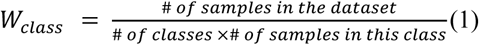

Training and hyper-parameter tuning were done through 5-fold stratified cross-validation. Hyper-parameters included a confidence score cutoff and an L2 regularization parameter. The confidence score cutoff was selected to classify the samples into positive and negative classes. We used the optimal cutoff to maximize the harmonic mean of sensitivity and specificity using equation (2) as suggested by Song et al.^20^:

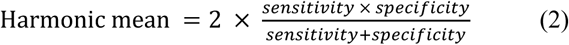

Also, the L2 regularization parameter was tuned by a log-spaced grid search in the cross-validation.

### Evaluation

The trained model was evaluated on our independent, held-out test dataset using standard machine learning performance metrics of the area under the ROC curve, F1 score, sensitivity, and specificity. Survival analysis was implemented with package ‘survival’^21^ in R version 3.3.3 (R Foundation for Statistical Computing, Vienna, Austria). Kaplan-Meier curves and log-rank tests were employed to examine the survival difference between the patients from different prediction, tumor grade, and treatment groups. For this analysis, patients from WHO 1973 grades 3 and 4 were combined into a single group due to small sample sizes. For further comparison with tumor grade, a multivariate Cox-proportional hazard model was also built with our model predictor and the WHO/ISUP classifications, while adjusting for the treatment information.

## Results

Our fully automated machine learning approach utilized variables with high reproducibility from a population-based dataset, without including less reliable measures like tumor grade to predict 10-year bladder cancer survival. In this evaluation, we applied our model on an independent test dataset (N = 286 patients). The final model achieved an area under the Receiver Operating Characteristic curve (AUC-ROC) of 0.77 (Figure 2), and an overall F1 score of 0.78 on the test dataset. Figure 2 shows that our final model reached a sensitivity of 0.65 (95% confidence interval 0.60 – 0.71) and a specificity of 0.79 (95% confidence interval 0.74 – 0.84). The accuracy of the model prediction was 0.76 on the testing dataset (95% confidence interval 0.71 – 0.81).

**Figure 2.**
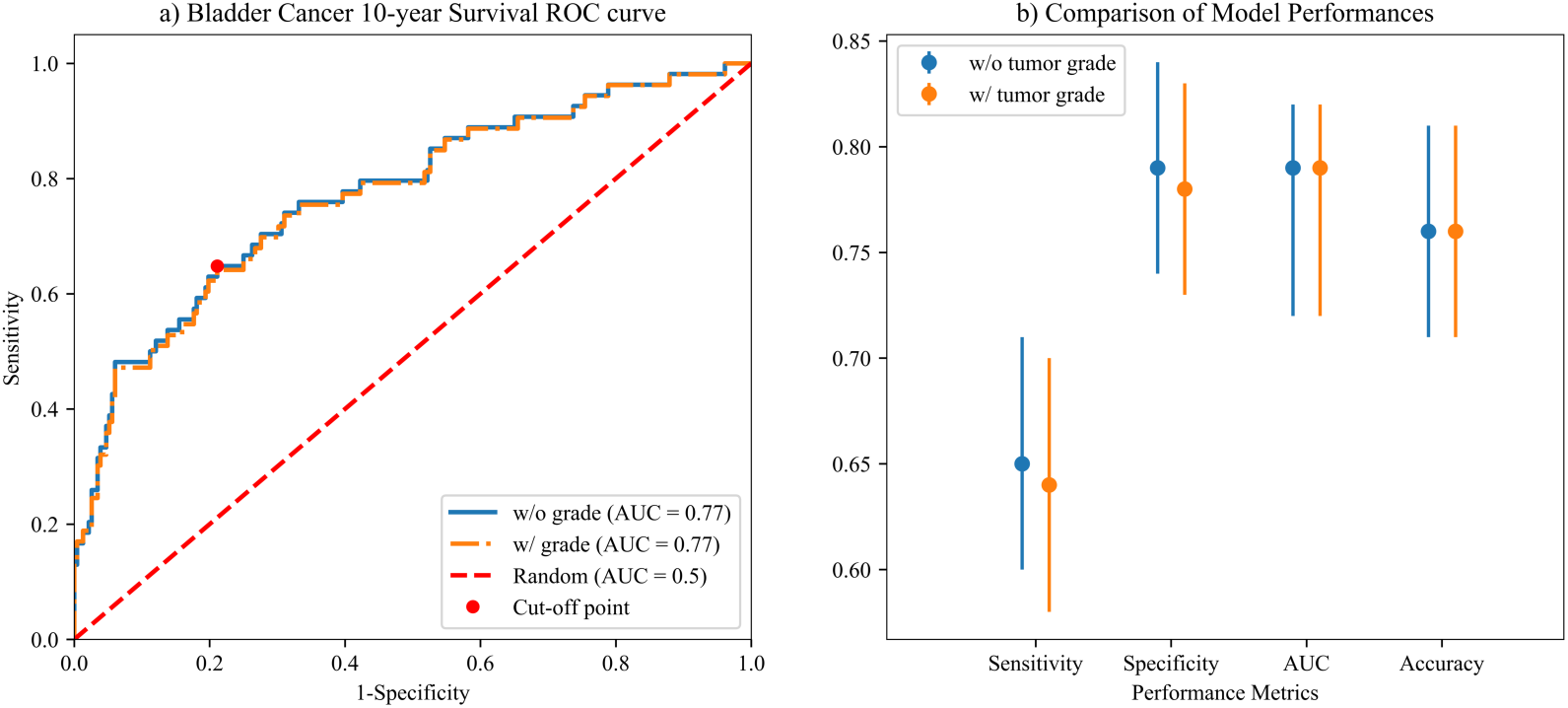
Model performance comparison of logistic regression model on the test dataset without and with WHO/ISUP tumor grade as a predictive feature. **a)** The receiver operating characteristic curve (ROC) of models’ predictions on all patients from the test dataset. The blue line shows the ROC of our final model (without tumor grade), and the orange line shows the ROC of the logistic model with tumor grade. The red dotted line represents the performance of a model with random predictions. The red dot represents the confidence score cut-off point of our final model to predict high and low-risk groups. **b)** Comparison of the model performance measurements with their 95% confidence intervals.

To show the effect and utility of tumor grade in our long-term prognosis predictions, a logistic regression model including WHO/ISUP tumor grade as an independent variable was trained on the same training dataset, using the previously described class weighting and hyper-parameter tuning approach. This model was tested for comparison with our final model that excluded the tumor grade. The model including tumor grade achieved same performances of in terms of AUC-ROC and F1 score, with slightly worse sensitivity and specificity, comparing to our final model (Figure 2). To show the directions and the magnitude of how each feature influences the prediction results, the coefficients of the predictive features of our final model are plotted in Figure 3.

**Figure 3.**
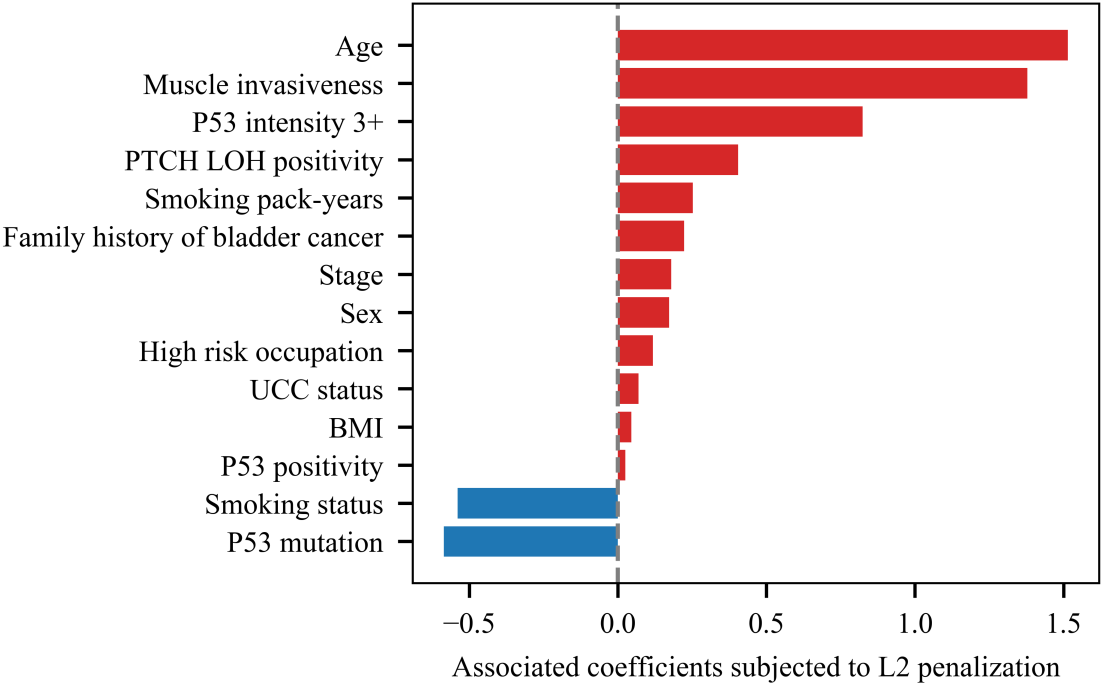
Coefficients of predictive features subjected to L2 penalization in final logistic regression model. The features with the highest impacts on making positive predictions (i.e., survival prognosis of fewer than 10 years from the initial diagnosis due to bladder cancer) included age, muscle invasiveness, and *P53* intensity.

Figure 4 depicts the Kaplan-Meier survival curve of Phase III patients stratified by our predicted risk groups, the existing tumor grading schemes, and patients’ treatment information. Our prediction model successfully distinguished the group with poorer prognoses from the rest, and the result was highly statistically significant (log-rank test p = 4.06 × 10^KL^). Of note, this result was more significant than both WHO 1973 (log-rank test p = 4.61 × 10^KM^) and WHO/ISUP grading schemes (log-rank test p = 5.99 × 10^KN^) for predicting long-term prognoses.

**Figure 4.**
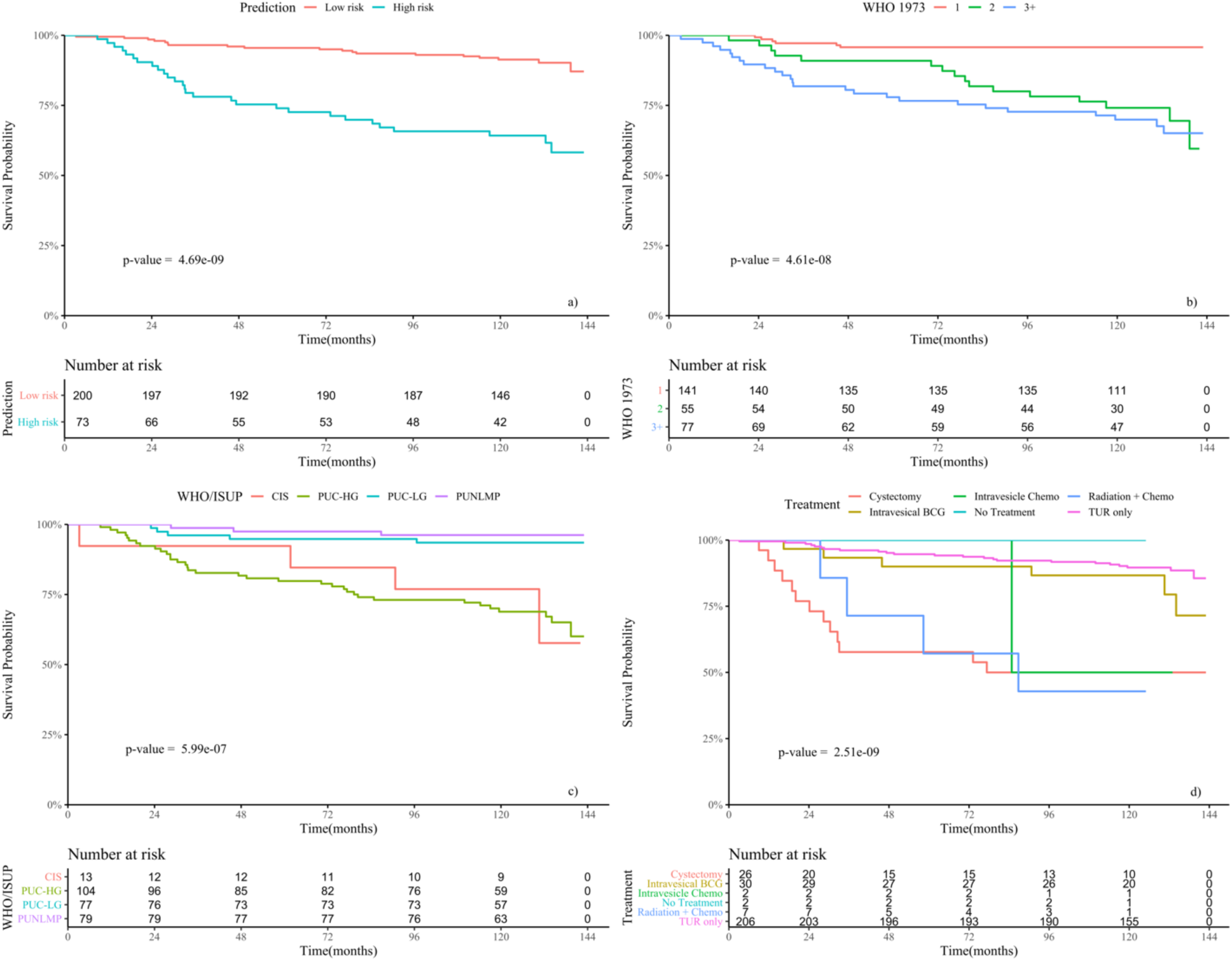
Survival analysis of bladder cancer patients from the test dataset stratified by **a)** our model’s predictions, **b)** the WHO 1973 classification, **c)** the WHO/ISUP classification, and **d)** treatment groups.

A multivariate Cox-proportional hazard model was built with our model predictor, the WHO/ISUP classification, and the patients’ first course of the treatment. As the result, our predictor achieved a hazard estimation of 2.99 (p = 0.0013, 95% confidence interval: 1.54 – 5.83) after adjustment for tumor grade based on the WHO/ISUP criteria or first course of treatment (Table 2). Although the magnitude of the effect was smaller than our model predictor, the treatment effect was statistically significant with respect to the 10-year survival, while WHO/ISUP is not (Table 2).

**Table 2.**
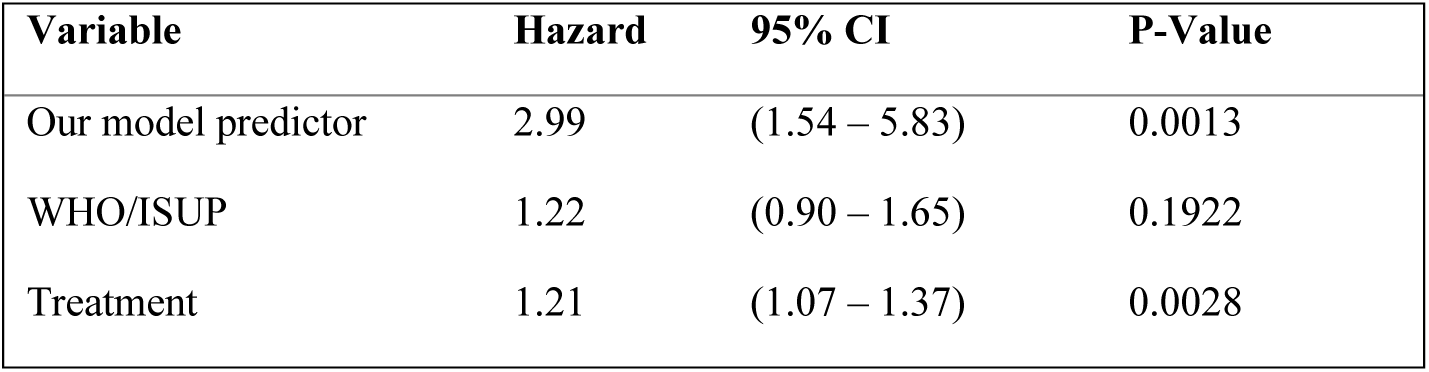
Result of multivariate Cox-proportional hazard model with our model predictor and the WHO/ISUP classification, adjusted for treatments.

| Variable | Hazard | 95% CI | P-Value |
| --- | --- | --- | --- |
| Our model predictor | 2.99 | (1.54 – 5.83) | 0.0013 |
| WHO/ISUP | 1.22 | (0.90 – 1.65) | 0.1922 |
| Treatment | 1.21 | (1.07 – 1.37) | 0.0028 |

## Discussion

In this study, we developed a machine learning model to predict 10-year survival of patients with bladder cancer. Our approach carries several unique advantages. First, we used a statewide, population-based dataset to train our model. The previous research efforts on predicting bladder cancer prognoses have relied on small datasets from single hospitals, which could be biased toward specific sub-populations^8^. Models from such studies need to be confirmed using data collected from multiple centers to ensure the generalizability of these models. Instead, our study design alleviated this problem of limited generalizability. Second, unlike previous studies, we excluded both the WHO 1973 and the WHO/ISUP classifications from the predictive features. These classifications exhibit high inconsistency based on clinicians’ personal experience, especially when resources of quality control are limited^22^. Third, our model also excluded treatment information to predict reliable prognoses before patients receive their first course of treatment and to serve as a constructive guide to aid treatment planning. The results from multivariate Cox regression analysis confirmed that our model’s predictions were strongly associated with patients’ survival. In addition, the effect of treatment was marginal and our predictor provided strong prognostic power regardless of the treatment information according to the multivariate Cox-proportional hazard model. As a result, our model achieved accurate performance on a held-out test dataset in predicting bladder cancer 10-year survival outcomes. Of note, exclusion of the tumor grade from the model did not penalize its performance. Instead, we observed a slight improvement in model’s sensitivity and specificity. This result suggests the potential for our approach to create a reliable tool, bypassing the inconsistent tumor grade, to support clinicians’ decision-making and patient management in clinical practice.

Our model prioritizes factors that are critical to bladder cancer prognosis to improve its prediction accuracy. For example, age, muscle invasiveness of the tumors, and smoking pack-years were among the highest-ranked variables in our final model. It is widely accepted that age is the most significant risk factor for bladder cancer occurrence^23^. Elderly patients have higher probabilities of developing muscle-invasive bladder cancer (MIBC), a form of bladder cancer with higher chance of metastasis and worse prognosis^1, 24^. MIBC patients are treated with more aggressive forms of treatment, such as radical cystectomy, which pose additional risks, complications, and side effects for less tolerant elderly patients^24^. Accumulation of environmental exposure to carcinogens, such as those from cigarette smoking, is also one of the major contributing factors to the incidence of bladder cancer^1, 25^.

An important advantage of our model is its ability to handle the imbalanced survival outcomes in the dataset. The estimated 10-year survival rate of bladder cancer patients is 70%, and patients with lower stage disease have much higher survival rates^1^. Thus, more patients have survived the 10-year mark than those who have not. A common weakness of using generic machine learning models on highly imbalanced data is the tendency for a bias toward the dominant class, which can result in low sensitivity. To minimize this bias, our model adopted a class weighting technique, placing greater emphasis on correctly predicting outcomes for the minor class. Additionally, we tuned the confidence score cutoff and chose a customized classification threshold that was well-suited for our imbalanced dataset. To further address this problem, we used the harmonic mean of sensitivity and specificity, and F1 score, rather than accuracy, as the primary evaluation metrics to account for the cost of both false positives and false negatives.

Lack of standardization for collected covariates has hindered many existing bladder cancer prognostic models from being adopted in clinical practice^26^. Our results in this study indicate that bladder cancer prognosis could benefit from the collection of several reproducible prognostic factors. Such information includes molecular features on *p53* alteration and refined demographical factors such as patients’ occupation and smoking in pack-years. *p53* alterations are highly prevalent among bladder tumors, and previous studies have implied the association between the mechanisms of *p53* alterations and the characteristics of bladder tumors^10, 27^. In addition, multiple other studies on developing tools for predicting bladder cancer prognostic outcomes also incorporated information regarding *p53* mutations, suggesting its potential value in clinical prognosis. While our automated machine learning pipeline is still applicable and beneficial for bladder cancer prognosis prediction in the presence of missing data, a complete collection of independent variables can achieve better accuracy for bladder cancer prognosis.

It is worth noting that the current approach has a few limitations. First, while our population-based dataset is more generalizable than a center-based dataset, the data collected from a state-wide registry may not represent a broader population, as the New Hampshire population is 92.3% White according to the United States Census Bureau in 2019^28^. Therefore, the presented model can become more clinically useful with the inclusion of more data, and further development and testing of the model. Another limitation of our prediction model is its relatively low sensitivity, even after the utilization of several machine learning improvement techniques to balance the decision boundary. This low sensitivity is likely inevitable in this study due to our skewed and imbalanced dataset with regard to the outcome variable. We plan to address this shortcoming in future work by collecting a more balanced dataset.

To improve the generalizability of our model, we plan to further validate our approach on a nationwide population dataset, such as the SEER database. Our next step is to enhance the model’s performance by training on more data points, and using data that is more complete. We plan to retrieve additional samples by identifying more recent cases from the New Hampshire Cancer Registry and collect more complete information on the relevant covariates as part of the ongoing New Hampshire Bladder Cancer Study. Of note, retraining the model on a larger, balanced dataset and doing further testing on the more recent data would be necessary before widely adopting it as a tool for statewide bladder cancer surveillance. Finally, we plan to include more discriminating factors in our predictive model. Such factors may include medical comorbidities retrieved from patients’ medical record, as well as additional clinicopathological features and genetic biomarkers. The latter data may not be routinely recorded in medical records, but it could be extracted from immunohistochemistry slides through automatic image analysis techniques using deep learning. We expect that integrating these clinicopathological features with the model presented here will improve its predictive performance.

